# Predicting Individual Variability in Task-Evoked Brain Activity in Schizophrenia

**DOI:** 10.1101/2020.05.25.114603

**Authors:** Niv Tik, Abigail Livny, Shachar Gal, Karny Gigi, Galia Tsarfaty, Mark Weiser, Ido Tavor

## Abstract

**BACKGROUND:** Patients suffering from schizophrenia demonstrate abnormal brain activity, as well as alterations in patterns of functional connectivity assessed by functional magnetic resonance imaging (fMRI). Previous studies in healthy participants suggest a strong association between resting-state functional connectivity and task-evoked brain activity that could be detected at an individual level, and show that brain activation in various tasks could be predicted from task-free fMRI scans. In the current study we aimed to predict brain activity in patients diagnosed with schizophrenia, using a prediction model based on healthy individuals exclusively. This offers novel insights regarding the interrelations between brain connectivity and activity in schizophrenia.

**METHODS:** We generated a prediction model using a group of 80 healthy controls that performed the well-validated N-back task, and used it to predict individual variability in task-evoked brain activation in 20 patients diagnosed with schizophrenia.

**RESULTS:** We demonstrated a successful prediction of individual variability in the task-evoked brain activation based on resting-state functional connectivity. The predictions were highly sensitive, reflected by high correlations between predicted and actual activation maps (*Median* = 0.589, *SD* = 0.193) and specific, evaluated by a Kolomogrov-Smirnov test (*D* = 0.25, *p* < 0.0001).

**CONCLUSIONS:** A Successful prediction of brain activity from resting-state functional connectivity highlights the strong coupling between the two. Moreover, our results support the notion that even though resting-state functional connectivity and task-evoked brain activity are frequently reported to be altered in schizophrenia, the relations between them remains unaffected. This may allow to generate task activity maps for clinical populations without the need the actually perform the task.

## Introduction

What goes wrong in a psychiatric patient’s brain that makes it so different from the healthy brain? This has been a challenging question for neuroscientists and psychiatrists for many years. Numerous studies have tried to answer this question regarding schizophrenia (SCZ), because of its extremely debilitating effects on the lives of patients and relatives, and its overwhelming economic burden on society (1–4). A Vast body of evidence from over 20 years of neuroimaging studies in schizophrenia suggests that one of its underlying causes is a disruption in brain connectivity (5,6).

The idea that schizophrenia is a disorder of brain connectivity had become widely accepted in the past two decades (6–8), mainly due to advances in neuroimaging that allowed investigation of in-vivo brain connectivity in the living human-brain (5). A Broadly used approach to measure brain connectivity non-invasively is to detect brain regions that show high temporal correlation in functional magnetic resonance imaging (fMRI) scans acquired at rest, i.e. when no explicit task is introduced (rs-fMRI). This method can be used to explore the architecture of functional brain networks, often referred to as resting-state networks (RSN’s) (9,10). Alterations in various RSN’s which are associated with high cognitive functions, such as the default-mode (11), salience (12) and fronto-parietal control (13) networks are consistently reported in schizophrenia, and are in line with the substantial cognitive deficit and behavioral changes that are among the hallmarks of this condition (1,14).

Alterations in task-associated brain function in schizophrenia are also very well documented. Task-fMRI studies use carefully designed behavioral tasks that participants perform in the scanner in order to localize functionally specialized brain regions that are active in response to cognitive demands. Task-induced activation is measured and localized by comparing the signal change between different cognitive states (9). Abnormal brain activation in many different fMRI tasks such as emotional response, decision making, response inhibition and working memory tasks was demonstrated in SCZ patients compared to healthy individuals (15,16).

Even though the abovementioned deficits in both functional connectivity and brain activity reported in schizophrenia may be in line with one another, they were traditionally studied mostly as two separate elements (9). In the last few years, studies have been trying to bridge this gap and establish the claim that brain activation while performing different tasks is closely related to brain connectivity. Such studies, focusing mainly on young healthy adults, have shown that functional connectivity closely corresponds with task-derived measures, and that RSNs qualitatively resemble task-evoked networks at the group level (17–19).

The fact that connectivity-derived brain networks correspond with task-fMRI activation maps formed the basis for the hypothesis that task-evoked brain activity could be predicted from task-free connectivity measures. Recently, by applying computational models to fMRI data, it has been demonstrated that functional connectivity measures obtained from scans acquired at rest can predict differences in fMRI activation across a range of cognitive paradigms (20). Using a simple model and carefully crafted predictors, it is possible to accurately map brain activity from rs-fMRI in individuals, even if their brain activation is unique or abnormal. This highlights a strong coupling between brain connectivity and activity that can be captured at the level of individual participants.

The abovementioned findings suggest that the relationship between functional connectivity and task-evoked brain activity in healthy individuals may be a constant intrinsic trait rather than a transient state (21). If it is in fact an intrinsic trait, it suggests that networks’ organization could be mapped directly to brain function. This means that given a specific set of connectivity measures it will be possible to predict brain activation patterns in individuals regardless of their specific brain attributes and even if they suffer from a pathology that effects these attributes.

The idea that task-evoked brain activity can be predicted from task-free functional connectivity in both healthy individuals and patients was supported by a study which demonstrated that brain activity could be successfully predicted in patients awaiting neurosurgery (22). However, these patients suffered from focal and well understood structural abnormalities. The question whether the close relationship between functional connectivity and brain activity is similar for individuals with brain pathologies that are more “diffuse” and “holistic", such as schizophrenia and other psychiatric disorders, remains unclear.

Successful predictions of brain activity from functional connectivity would allow studying challenging populations that are usually not compliant with task performance, and therefore may be a promising method to study brain function in psychiatric patients (23). Moreover, reducing fMRI protocol to a single rs-fMRI scan, instead of a series of demanding cognitive tasks, would save a substantial amount of resources and diminish scan related inconvenience. But above all, this approach has the potential to deepen our understanding of how brain connectivity and activity are interrelated in the pathological brain and may further the understanding of the underlying mechanisms of schizophrenia.

In the current study we aimed to investigate the relationship between functional connectivity and task-evoked brain activity in SCZ patients by predicting task-evoked brain activity from task-free functional connectivity measures. In order to achieve that we trained a prediction model based on functional connectivity measures extracted from task-free scans of healthy controls and applied the model to predict working memory task-evoked brain activity in SCZ patients. We hypothesized that such predictions will be successful, in terms of both sensitivity, represented by high overlap between predicted and actual brain activation maps, and specificity, represented by accurate prediction of individual-unique activation patterns.

## Materials and Methods

### Participants

The dataset was acquired in Sheba Medical Center, Ramat-Gan, Israel. It originally consisted of 112 participants: 89 healthy volunteers and 23 patients diagnosed with schizophrenia (SCZ). All participants were 18-55 years old Hebrew speakers and had no history of neurological conditions. The control group had no history of psychiatric conditions. The research protocol was approved by the Institutional Review Board of Sheba Medical Center. All participants signed an informed consent form. Nine healthy volunteers and three SCZ patients were excluded from the study due to insufficient task performance or the lack of vital MRI scans. The final dataset used for analysis consisted of 80 healthy volunteers and 20 patients. Clinical and demographic characteristics are described in table 1. Individual level characteristics for SCZ group are described in table S1.

**Table 1.** Clinical and demographic characteristics. SCZ=Patients diagnosed with schizophrenia.

|  | Control (N=80) | SCZ (N=20) |
| --- | --- | --- |
| **Age (years)**<br>Mean (STD) | 35.42 (11.97) | 29.3 (6.3) |
| **Gender**<br>% Male | 56.25 | 80 |
| **PANSS total**<br>Mean (STD) | - | 50.64 (12.83) |
| **PANSS general**<br>Mean (STD) |  | 23.47(5.47) |
| **PANSS positive**<br>Mean (STD) | - | 11 (4.99) |
| **PANSS negative**<br>Mean (STD) | - | 16.18 (5.16) |

### Clinical Assessment

All participants in the SCZ group met the criteria for schizophrenia according to DSM-5 and were diagnosed by a psychiatrist. Symptoms severity was evaluated by using the Positive and Negative Syndrome Scale (PANSS) for schizophrenia (24). Out of 20 Patients used for the analysis, 17 had available PANSS scores. Averages of these scores are described in table 1 and individual scores in table S1.

### fMRI Working Memory Task

All participants conducted the widely used fMRI working memory (WM) N-back task (25,26). Differences between control participants and SCZ patients in this task have been reported consistently, in both brain activation patterns and task performance (11,27,28). A detailed description of the task design can be found in the supplemental information.

### MRI Acquisition

Participants underwent an MRI session which included anatomical, task-fMRI and rs-fMRI scans. Scans were acquired on a 3 Tesla whole body MRI system (GE Signa HDxt, version 16 VO2) equipped with an eight-channel head coil.

Anatomical high-resolution (1mm^3^, Matrix size 256 × 256, FOV 25.6 cm) images of the entire brain were acquired, using a standard 3D inversion recovery prepared fast spoiled gradient echo pulse (FSPGR) T1 weighted sequence. Additional anatomical sequences (T2w and fluid-attenuated inversion recovery (FLAIR)) were acquired for radiological screening.

Working memory task functional scans were acquired with a T2*-weighted gradient-echo echo-planar protocol (GE-EPI) using the following parameters: TR=3s; TE=30-35ms; matrix size 64×64, FOV 22×22 cm, and up to 40 contiguous oblique axial slices covering the whole brain. The resulting voxel size was 3.4 mm^3^.

Rs-fMRI protocol was almost similar to the task fMRI protocol, except that TR was reduced to 2s and scan time was 9:53 minutes. Participants were instructed to close their eyes during the scan.

### fMRI Preprocessing and Individual Statistics

fMRI preprocessing (see Figure 1) for both rs-fMRI and N-back task was carried out using FMRIB Software Library (FSL v5.0.10) (29) and included high-pass filtering at 0.01Hz, correction for motion artifacts, linear registration to the T1w anatomical scan, nonlinear registration to 152MNI space, and smoothing with 5mm gaussian kernel. Residual noise was cleaned using FMRIB’s ICA-based Xnoiseifier (FIX) (30), which is a semi-automatic ICA based method to identify and remove structured noise from fMRI scans. Then each scan was resampled onto the set of 91,282 “grayordinates” (31) in standard space which was used for surface representation using the HCP’s Connectome Workbench visualization and discovery tool (32). Individual level statistical analysis to generate task-evoked activation maps for each participant was performed using the FEAT pipeline in FSL (33). Task activation maps were generated for the 2back>0back contrast representing brain activity associated with working memory (25).

**Figure 1.**
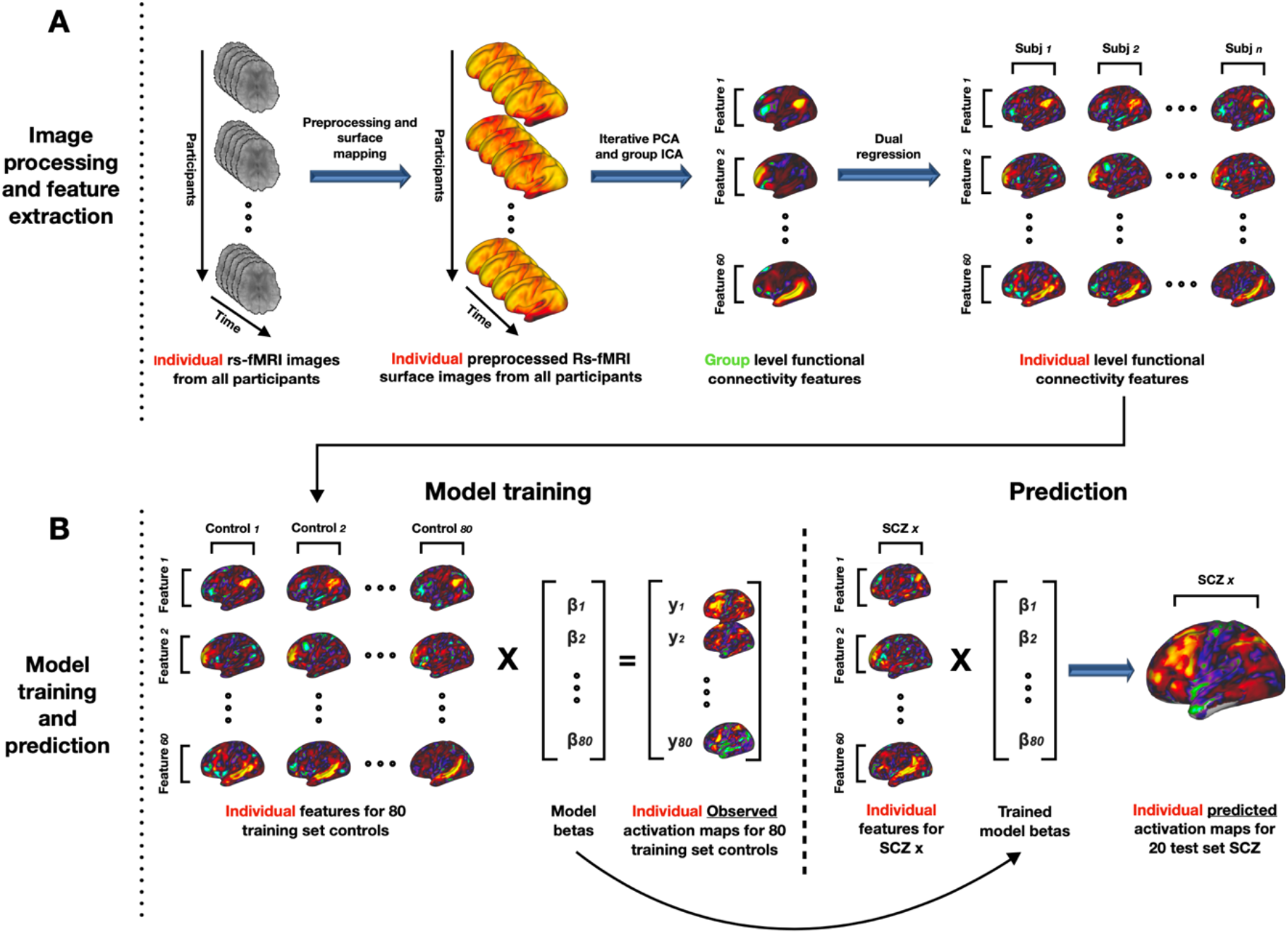
Pipeline for task-evoked brain activity prediction from task-free functional connectivity measures. fMRI preprocessing and feature extraction (A) included standard preprocessing for resting-state functional MRI (rs-fMRI) scans using the FMRIB Software Library (FSL) and mapping into 91282 “Grayordinates” surface. Then, iterative principal component analysis (PCA) followed by group independent component analysis (ICA) were performed to yield group level functional connectivity maps (features). Next, dual regression was applied to generate individual features for each participant. This process was followed by the model training and prediction pipeline (B). Our GLM based prediction model was trained on features extracted solely from healthy controls (training set) and validated using a leave-one-out routine. Last, the trained model was applied on the SCZ patients (test set) and yielded a predicted activation map for each participant.

### fMRI group analysis

As mentioned above, group differences in brain activity between healthy and SCZ participants in the N-back task are well documented (11,27,28). However, it is important to validate that such differences are found in our specific dataset in order to ensure that successful prediction is not simply a result of no variability between the groups. In order to test for group differences in task evoked brain activation, group level analysis was performed using FSL’s FEAT pipeline. Z-score activation maps depicting group differences in brain activation were created using FMRIB’s local analysis of mixed effects (FLAME) (34). The activation maps were then thresholded using a Gaussian-two-Gammas mixture model (35), where the Gaussian represents the noise and the two Gamma distributions represent positive and negative activations. The positive and negative thresholds are chosen to correspond to the medians of the two Gamma distributions.

### Functional connectivity-based classification

It is well established that SCZ patients demonstrate altered patterns of brain connectivity compared to controls (5,6,11). Before exploring the predictability of task-evoked activity from connectivity measures, we aimed to assert these differences in our dataset. In order to achieve this, we trained a classifier to distinguish between controls and SCZ based on functional connectivity measures.

Feature extraction for classification was conducted by averaging the preprocessed rs-fMRI time courses within each of 100 cortical parcels, where each parcel was assigned to one of seven brain networks, according to a parcellation by Schaefer et al. (36). Pearson’s correlation coefficient was calculated for each dyad of parcels, resulting in 4950 correlation scores, which were used as features for the classification model. All features were normalized by removing the mean and scaling to unit variance.

Then, we utilized an elastic-net logistic regression model, that combines L1 and L2 penalties for regularization, in order to classify each participant to one of our two groups. The L1/L2 ratio (α) and the regularization factor (λ) were chosen in a stratified 5-fold cross-validation procedure. In order to determine chance level rates and utilize them to test for statistical significance we used a 5000 iterations permutation test in which our classification labels (control or SCZ) were shuffled randomly. We calculated 4 scores to determine prediction success: AUC (Area under the receiver operating characteristics curve), accuracy, sensitivity and specificity. For more details see the supplemental information.

### Prediction of task activity from functional connectivity measures

The prediction pipeline was adapted from Tavor et al., 2016 (20). We used data from 80 control participants as a training set. Feature extraction included dimensionality reduction of preprocessed rs-fMRI maps by iterative group principal component analysis (PCA) (37), yielding 200 group level principal components. Next, group-level spatial independent component analysis (ICA) (38) was carried out on cortical data using fast ICA (39) to define a set of seeds for 60 cortical functional connectivity maps. Then dual regression (40) was performed on the group level connectivity maps to create individual-level functional connectivity maps that were used as features for the prediction model. More details regarding the feature extraction process can be found in figure 1. and the supplemental information.

A linear model was used to map the functional connectivity features to the individual task-evoked activation maps. The model was first trained on the 80 healthy controls training set. Regression coefficients (betas) were calculated for each participant, and then averaged for *n* − 1 participants each time to execute a leave-one-out (LOO) prediction routine and generate a predicted task activation map for each control participant. Then, the trained model was utilized to predict task activation maps in 20 SCZ patients that were kept out in previous steps.

## Results

### N-back working memory task group differences

Group differences between SCZ patients and healthy controls in the 2back>0back contrast of the N-back task were tested. As expected, significant differences in activation were found in various cortical regions. Healthy controls displayed higher activation mainly in areas that correspond with the frontoparietal control network (41) such as the dorsolateral prefrontal cortex (DLPFC), medial frontal cortex, inferior parietal lobule and posterior temporal regions. SCZ patients displayed significantly higher activations mostly in occipital visual areas and in the left insula. These group differences are presented in figure 2.

**Figure 2.**
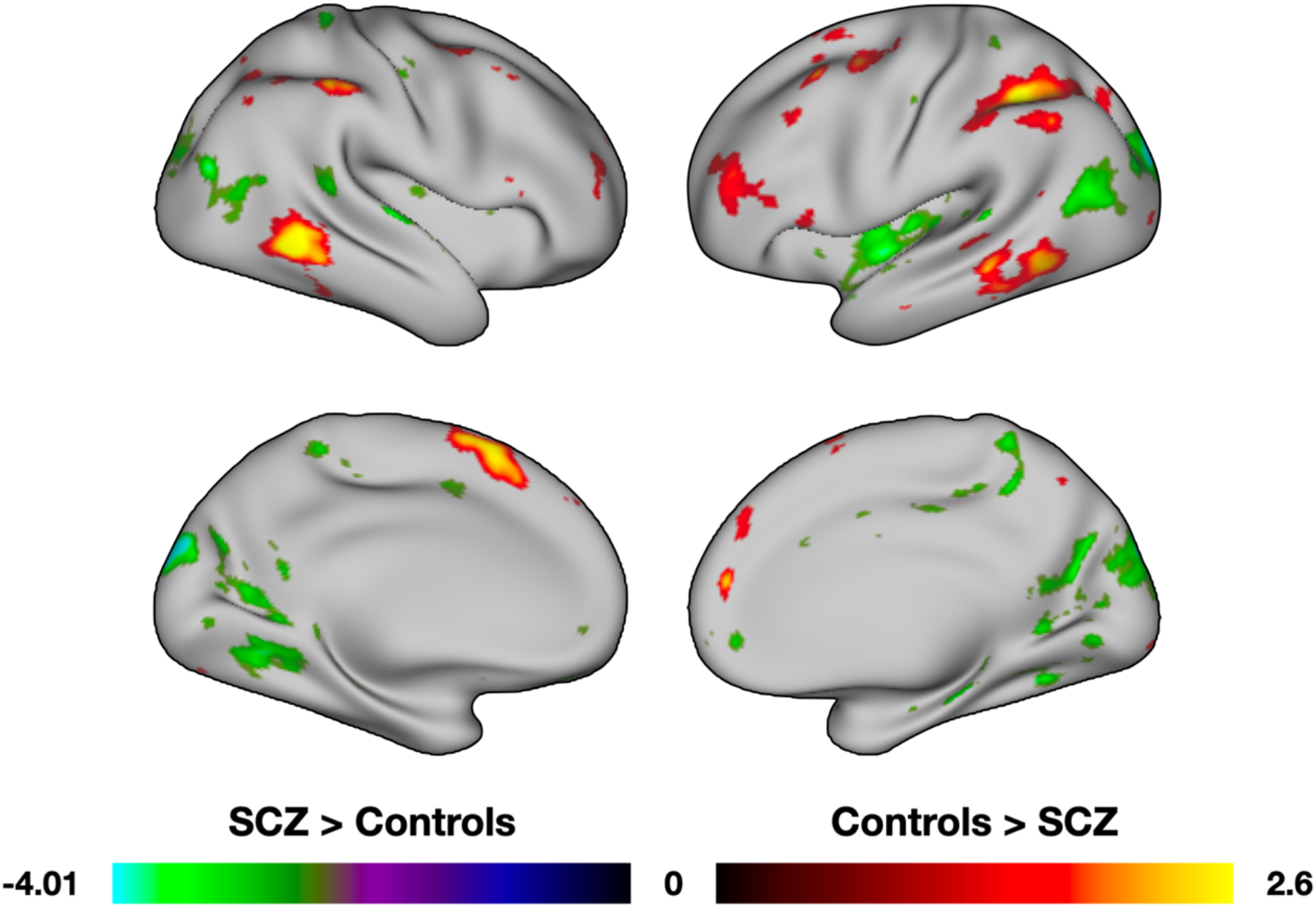
Brain activation maps showing group differences between control and SCZ in the N-back working memory 2back>0back contrast. The maps are thresholded using Gaussian-two-Gammas mixture model. The positive threshold was set as Z=1.08 and the negative as Z=−2.41. Areas that had higher activation in controls are displayed in hot colors and areas that had higher activation in SCZ are displayed in cold colors.

### Functional connectivity-based classification

We used an elastic-net logistic regression classifier to explore the ability to successfully classify our participants as either control or SCZ using rs-fMRI derived functional connectivity measures. We used a 5-fold cross-validation routine to choose the optimal L1/L2 ratio (α = 0.7) and the regularization factor (λ = 0.77). In order to test the statistical significance of our classification we ran a permutation test and used it to compute chance level rates for 4 classification performance scores. All scores were found significant compared to the computed chance level: *AUC* = 0.85, (*p* = 0.0002), *Accuracy* = 0.78, (*p* = 0.003), *Sensitivity* = 0.8, (*p* = 0.003) and *Specificity* = 0.77, (*p* = 0.0074). These results indicate successful classification of participants into control and SCZ using functional connectivity features (Figure 3A).

**Figure 3.**
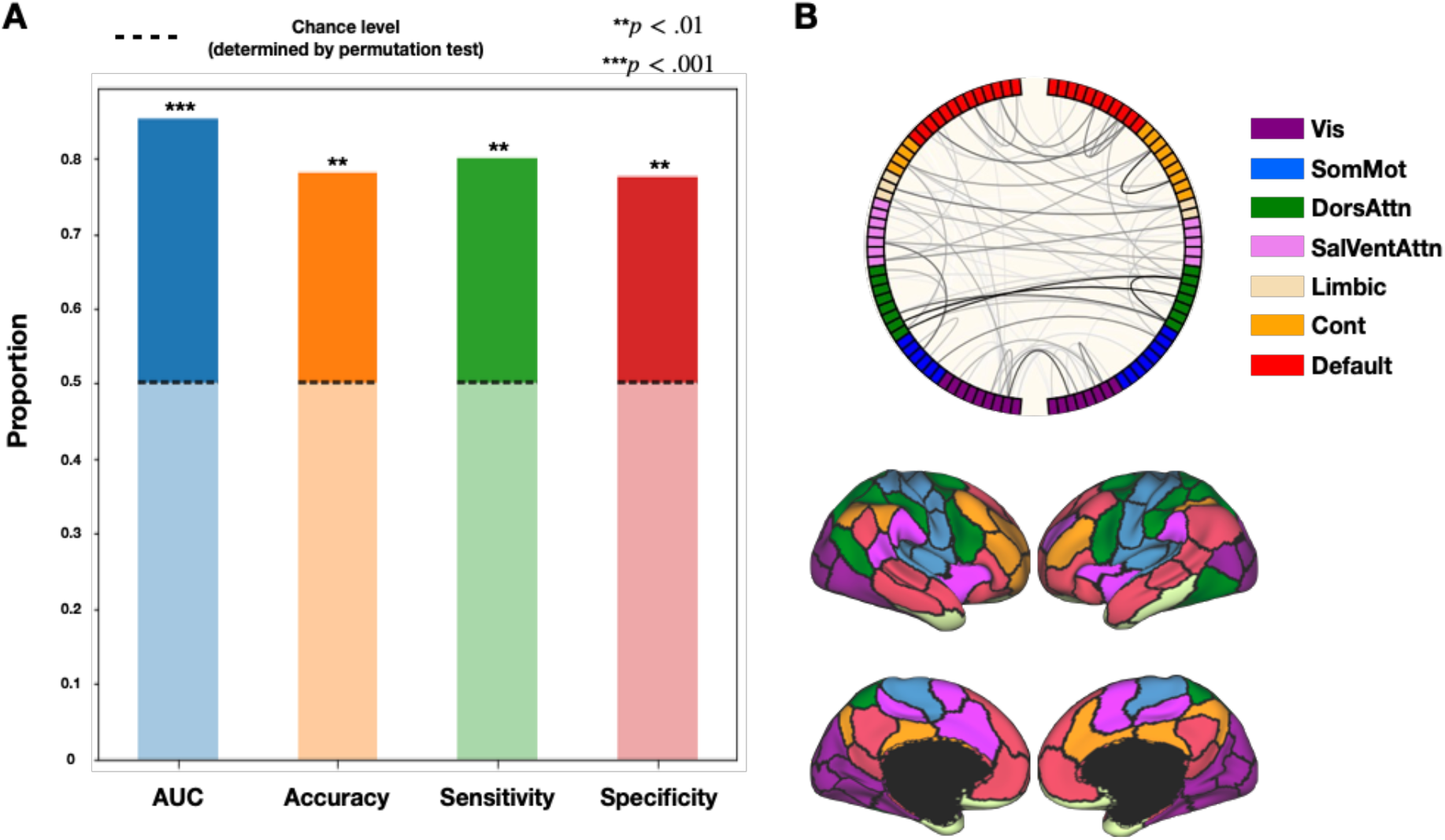
Classification of participants into control and schizophrenia groups. (A) shows the results of a permutation test designed to determine classification success, Evaluated by, 4 scores: area under the ROC curve (AUC), accuracy, sensitivity and specificity. Computed chance level for each score is marked by a dashed line on the bars. (B) shows the 100 edges that contributed most to the classification plotted on a circular graph. Each edge in the graph is one of the features used for classification, and its importance is depicted by the color in grayscale (i.e., darkest edges contributed most). Each half of the plot represents one hemisphere. The nodes are colored according to the parcellation by Schaefer et al., (2018) also shown in (B).

To explore which features were most relevant to the classification, we sorted the features by absolute classification beta values and extracted the top 100 features. Out of these features, 55 were inter-hemispheric connections and 45 intra-hemispheric: 26 in the left hemisphere and 19 in the right. Out of the 56 inter-hemispheric connections, 31 were homotopic, meaning they connected nodes that are assigned to the same network in different hemispheres. Therefore, the proportion of homotopic connections in the top 100 contributing features was 31% and 70% in the top 10, even though they amount to only 8% of the total features used for classification. Figure 3B presents the top 100 contributing features on a circular graph. Each edge in the graph is a features used for the classification, and its importance is depicted by the color in grayscale. Each node is colored by networks according to the parcellation by Schaefer et al. (36) (Materials & Methods, Figure 3B). To further explore the important role homotopic connections, have in differentiating SCZ patients from healthy controls we repeated the classification using only homotopic connections as features. This process improved our classification ability even more:: *AUC* = 0.89, (*p* = 0.0002), *Accuracy* = 0.82, (*p* = 0.008), *Sensitivity* = 0.8, ( *p* = 0.0038), *Specificity* = 0.825, (*p* = 0.0018) (see figure S4 and Table S2).

### Brain activity prediction

Our GLM-based prediction model was trained on 80 healthy controls. Using a Leave-one-out routine, a predicted cortical activation map was created for each participant in the training set. Then the trained model was used to predict cortical activation maps in the test set that consisted of 20 SCZ patients. Exemplar maps showing the predicted and actual activations and the substantial overlap between them in both control and SCZ groups can be found in figure 4A.

**Figure 4.**
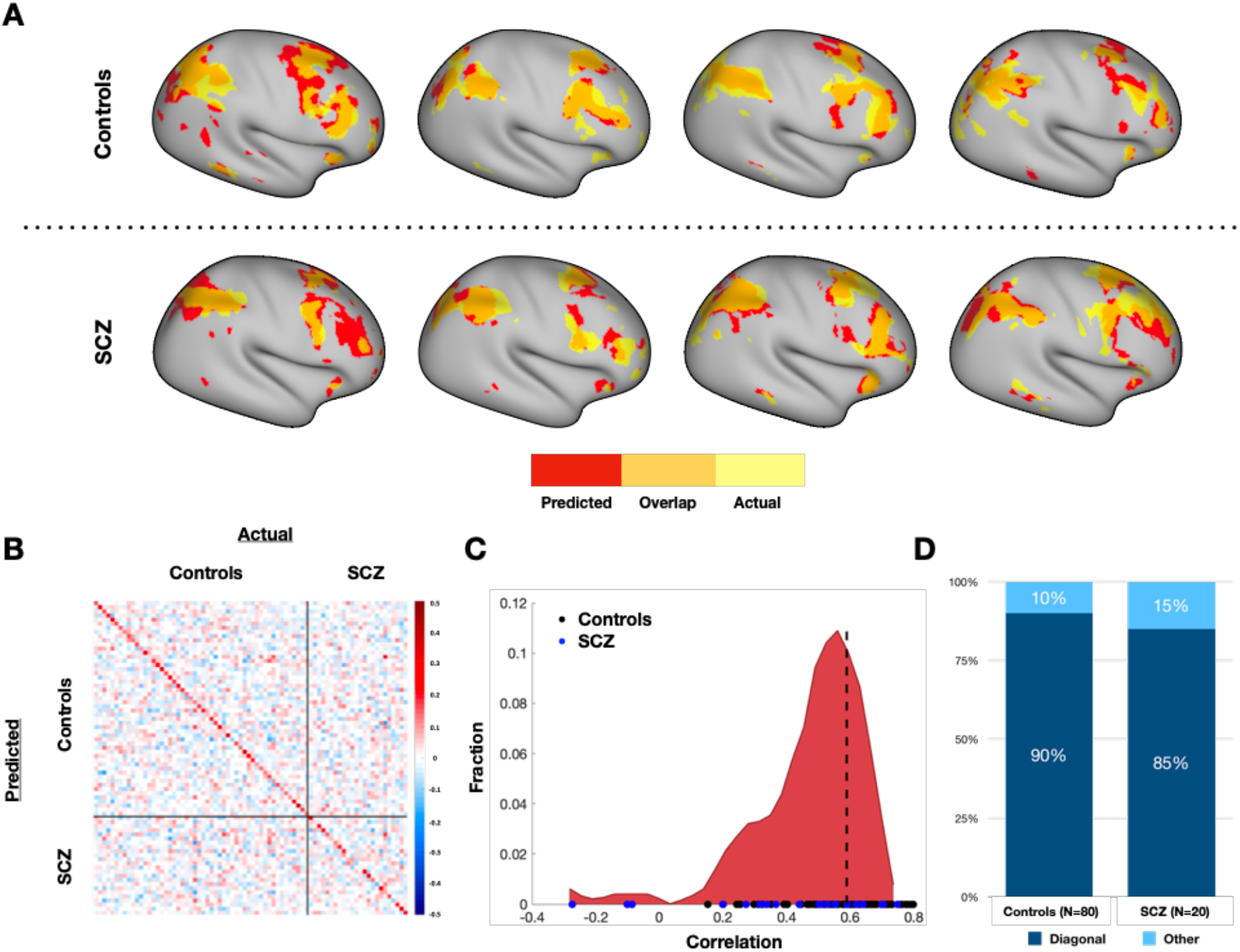
N-back task-evoked brain activity prediction. (A) shows thresholded binary predicted (red) and actual (yellow) task activation maps and the substantial overlap between them (orange) for 4 control and 4 SCZ participants. (B) shows a correlation matrix between predicted and actual activation maps in both groups. Rows and columns are normalized by removing the mean in order to account for higher variability in actual than predicted maps. The diagonal represents correlations between predicted and actual maps in the same individuals, hence the diagonality of the matrix indicates high prediction specificity. (C) shows a histogram of the off-diagonal values. The markers on the x-axis are the diagonal values, blue for the SCZ group and black for the control group. The dashed line indicates the median of diagonal correlations. (D) shows the proportion of participants in each group that the diagonal (individual-specific) prediction was the most accurate one for them.

We calculated the correlations between the actual and predicted activation maps of all pairs of participants (Figure 4B). The diagonal of the resulting correlation matrix represents correlations between the predicted and actual activation maps of the same participant (diagonal correlations). The rest of the matrix represents correlations between predicted and actual activation maps of different participants (off-diagonal correlations). An accuracy score was calculated as the average Pearson correlation between participants’ actual and predicted activation maps (*Mean* = 0.544, *Median* = 0.589, *SD* = 0.193). The specificity score was defined as the diagonality index, calculated as the average difference between diagonal and off-diagonal correlations (*DI* = 0.063). Diagonality of the correlation matrix that indicates high prediction specificity was quantitatively verified by a Kolmogorov-Smirnov test between diagonal and off-diagonal correlations, in which diagonal correlations were found significantly higher (*D* = 0.25, *p* < 0.0001). A histogram of diagonal and off-diagonal values is shown in Figure 4C. In order to further quantify prediction specificity, we calculated the proportion of participants for whom the diagonal correlation was highest, i.e. their actual activation map resembled their own predicted map more than any other predicted map. The diagonal correlation was highest for 90% (72 out of 80) of the control group and 85% (17 out of 20) of the SCZ group (Figure 4D).

To determine which features contributed most to the prediction, the regression coefficient (beta) value of each feature was averaged across all training set participants. The features were then sorted by absolute beta value. A bar plot of average beta values and the top 5 contributing features is shown in Figure 5.

**Figure 5.**
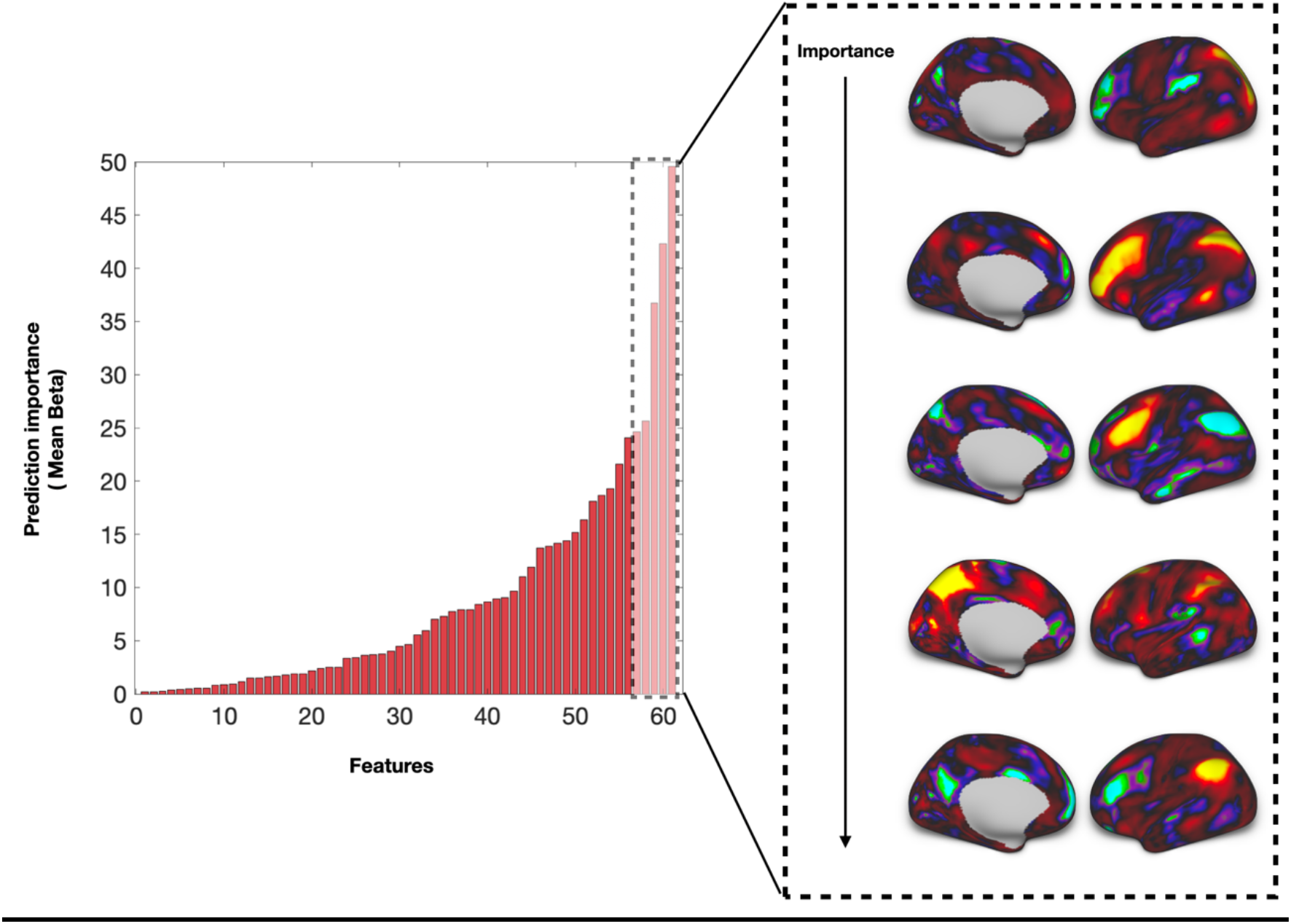
Features (connectivity maps) that were used for prediction of N-back task-evoked brain activity, sorted by their importance for prediction (determined by absolute GLM regression coefficient value). Surface representation of the top 5 contributing features is also shown.

In order to make sure that our ability to predict cortical activations does not depend on symptoms severity we calculated the correlation between prediction sensitivity (diagonal correlations) and PANSS scores for the 17 SCZ patients for which we had valid documented PANSS scores. We calculated correlations with positive symptoms (*r* = −0.13, *p* = 0.61), negative symptoms (*r* = 0.23, *p* = 0.37), general pathology score (*r* = 0.2, *p* = 0.44) and total PANSS score (*r* = 0.12, *p* = 0.63). All correlations were found statistically insignificant (using *P* < 0.05).

## Discussion

In the current study we provide the first evidence of successful predictions of fMRI task-evoked brain activity in patients diagnosed with schizophrenia. Moreover, the predictions were accomplished using a model trained on a group of healthy controls exclusively, which supports our hypothesis that the relationship between functional connectivity and task-evoked brain activity is a stable intrinsic trait. Hence, our findings suggest that the functional organization of brain networks inferred from task-free fMRI scans can be mapped directly to task-evoked brain function. We demonstrate that given a specific set of connectivity measures, it is possible to predict brain activation patterns in individual participants, regardless of individual-specific brain attributes such as a diagnosed brain pathology.

Our findings are in line with previous literature describing schizophrenia as a disorder of brain connectivity (5,6,11). Along with the growing body of evidence linking connectivity directly to task-evoked brain activity (17,18,20,42) and cognitive function (13,43), the current study provides support to the claim that abnormal brain activation patterns and altered cognitive function, which are widely reported in schizophrenia, may correspond directly to changes in the architecture of connectivity-derived brain networks. The successful predictions of task-evoked brain activity from task-free functional connectivity measures suggest that the abnormalities observed in SCZ in task-performance and the related brain activity are manifested at the level of functional connectivity and may even derive directly from connectivity alterations. Therefore, it is important to deepen our understanding of the mechanism underlying these connectivity alterations, that may be a potential target for developing biomarkers or disease modifying interventions (23).

Importantly, we demonstrated predictions that are both sensitive and specific. This is a core issue when trying to make individual predictions because it is cardinal to not only predict the shared variance across all individuals, but also account for the variance that is unique for every individual. Modern “dimensional” approaches, such as the Research Domain Criteria approach spearheaded by the NIMH (44), calls to move from statistically based dichotomic diagnoses towards describing individual specific phenotypes as a continuum. Understanding that these approaches will become more and more prominent in psychiatric research, we must be able to draw individual-level conclusions that will ultimately not rely on dichotomic diagnosis (21). The current study goes hand in hand with these approaches, as our findings suggest a framework in which we rely upon learning the unique relationship between an individual’s connectivity and brain activation patterns and therefore do not have to train the prediction model on diagnosed patients in order to predict their brain activation patterns. Hence, our ability to make such predictions will not be affected if the criteria for diagnosis change dramatically.

We examined the differences between the SCZ and control groups in both functional connectivity and task-evoked brain activity, in order to confirm that our predictive ability could not be explained by the lack of variability between the groups. Previous studies have shown that SCZ patients and healthy controls can be successfully differentiated using machine learning based classifiers utilizing features derived from functional connectivity (45–47). Consistent with these findings we showed that the healthy controls and SCZ patients in the current dataset can be accurately differentiated based on functional connectivity measures.

When examining feature importance, we noticed that homotopic connections, meaning connections between areas within the same network in both hemispheres, are cardinal for classification. As evidence, their proportion in the top 100 important connections is 31%, and 70% of the top 10, while their proportion from the total features is only 8%. Moreover, when we used only homotopic connections as features, our classification ability improved. These findings provide support to claims made by previous studies that demonstrated reduction in homotopic functional connectivity in SCZ patients (48,49), and even in their siblings (50), and therefore highlights the need to better understand alterations in homotopic connectivity in schizophrenia.

As for the task-evoked brain activity results, we found differences between SCZ patients and healthy controls in the N-back working memory task. Such differences were also reported in previous studies (11,27,28). Even though task fMRI findings in schizophrenia seem to be rather inconsistent, a large body of evidence has reported a reduction of brain activity in areas related to the fronto-parietal control network (41), such as the DLPFC and the inferior parietal lobule, when working memory is a major factor in the performed task (51). These are brain regions that are highly associated with working memory, thus it is not surprising that patients demonstrated reduced activity in these regions. When examining the features that contributed most to task prediction, many of them are also associated with the fronto-parietal control network, which is also in line with the major role this network plays in working memory and other high cognitive functions (52).

There are a few methodological considerations that needs to be taken into account when interpreting our results. The first is related to the quality of the imaging data. While originally the predictability of brain activity from connectivity was demonstrated on very high quality datasets (provided by the HCP) (20), the current dataset did not meet the same standard, as often happens with clinical data. However, the fact that we were still able to show accurate predictions emphasizes that the relations between brain connectivity and activity are very robust and could therefore be detected even with sub-optimal data. Nevertheless, it might be possible to get even more accurate results with a larger higher-quality dataset. Another issue is one that almost all fMRI schizophrenia studies suffer from: the fact that we use long MRI protocols and a demanding cognitive task creates a bias towards high functioning patients that might not be a good representation of the population (23). To validate that this issue is not cardinal in our study, we showed that our ability to make individual-specific predictions does not depend on symptoms severity scores (PANSS). Finally, it is important to generalize our findings beyond schizophrenia to other psychiatric conditions. To achieve this, a large dataset of various fMRI tasks performed by patients diagnosed with various conditions should be collected.

In conclusion, we demonstrate a framework to predict fMRI task-evoked brain activity in SCZ using a training set consisting of healthy controls exclusively. This may be a promising approach for studying connectivity and brain function in SCZ, as it allows drawing conclusions that typically require hours of tedious scanning and task performance, with only one short MRI scan at rest. Moreover, our results support the notion that interrelations between functional connectivity and brain activity do not depend on volatile factors. Rather, they may be an intrinsic trait underlying the functional and the resulting behavioral abnormalities in SCZ. Therefore, future studies should not consider brain activity and connectivity as two different elements but try to understand how they integrate and interact in the healthy as well as the pathological brain.

## Acknowledgements and disclosures

The authors acknowledge with thanks the support of the Israel Science Foundation (ISF grant no. 1603/18). The authors report no biomedical financial interests or potential conflicts of interest.

## Notes

### Competing Interest Statement

The authors have declared no competing interest.

## References

1. Heinrichs RW, Zakzanis KK (1998): Neurocognitive deficit in schizophrenia: A quantitative review of the evidence. Neuropsychology 12: 426–445.

2. McCutcheon RA, Reis Marques T, Howes OD (2020): Schizophrenia - An Overview. JAMA Psychiatry 77: 201–210.

3. Cloutier M, Aigbogun MS, Guerin A, Nitulescu R, Ramanakumar A V., Kamat SA, et al. (2016): The economic burden of schizophrenia in the United States in 2013. J Clin Psychiatry 77: 764–771.

4. Velthorst E, Reichenberg A, Kapra O, Goldberg S, Fromer M, Fruchter E, et al. (2016): Developmental trajectories of impaired community functioning in schizophrenia. JAMA Psychiatry 73: 48–55.

5. Van Den Heuvel MP, Fornito A (2014): Brain networks in schizophrenia. Neuropsychol Rev 24: 32–48.

6. Lynall M-E, Bassett DS, Kerwin R, McKenna PJ, Kitzbichler M, Muller U, Bullmore E (2010): Functional Connectivity and Brain Networks in Schizophrenia. J Neurosci 30: 9477–9487.

7. Dong D, Wang Y, Chang X, Luo C, Yao D (2018): Dysfunction of Large-Scale Brain Networks in Schizophrenia: A Meta-analysis of Resting-State Functional Connectivity. Schizophr Bull 44: 168–181.

8. Camchong J, MacDonald AW, Bell C, Mueller BA, Lim KO (2011): Altered functional and anatomical connectivity in schizophrenia. Schizophr Bull 37: 640–650.

9. Bijsterbosch J, Smith SM, Beckmann CF (2017): Introduction to Resting State fMRI Functional Connectivity. Oxford Neuroimaging Primers.

10. van den Heuvel MP, Hulshoff Pol HE (2010): Exploring the brain network: a review on resting-state fMRI functional connectivity. Eur Neuropsychopharmacol 20: 519–34.

11. Whitfield-Gabrieli S, Thermenos HW, Milanovic S, Tsuang MT, Faraone S V., McCarley RW, et al. (2009): Hyperactivity and hyperconnectivity of the default network in schizophrenia and in first-degree relatives of persons with schizophrenia. Proc Natl Acad Sci U S A 106: 1279–84.

12. Palaniyappan L, Simmonite M, White TP, Liddle EB, Liddle PF (2013): Neural primacy of the salience processing system in schizophrenia. Neuron 79: 814–828.

13. Godwin D, Ji A, Kandala S, Mamah D (2017): Functional Connectivity of Cognitive Brain Networks in Schizophrenia during a Working Memory Task. Front psychiatry 8: 294.

14. Knowles EEM, Weiser M, David AS, Glahn DC, Davidson M, Reichenberg A (2015): The Puzzle of Processing Speed, Memory, and Executive Function Impairments in Schizophrenia: Fitting the Pieces Together. Biol Psychiatry. https://doi.org/10.1016/j.biopsych.2015.01.018

15. Minzenberg MJ, Laird AR, Thelen S, Carter CS, Glahn DC (2009): Meta-analysis of 41 functional neuroimaging studies of executive function in schizophrenia. Arch Gen Psychiatry 66: 811–822.

16. Gur RE, McGrath C, Chan RM, Schroeder L, Turner T, Turetsky BI, et al. (2002): An fMRI study of facial emotion processing in patients with schizophrenia. Am J Psychiatry 159: 1992–1999.

17. Smith SMM, Fox PMMPTT, Miller KLL, Glahn DCC, Fox PMMPTT, Mackay CEE, et al. (2009): Correspondence of the brain’s functional architecture during activation and rest. Proc Natl Acad Sci 106: 13040–13045.

18. Cole MW, Bassett DS, Power JD, Braver TS, Petersen SE (2014): Intrinsic and task-evoked network architectures of the human brain. Neuron 83: 238–251.

19. Krienen FM, Thomas Yeo BT, Buckner RL (2014): Reconfigurable task-dependent functional coupling modes cluster around a core functional architecture. Philos Trans R Soc B Biol Sci 369. https://doi.org/10.1098/rstb.2013.0526

20. Tavor I, Jones OP, Mars RB, Smith SM, Behrens TE, Jbabdi S (2016): Task-free MRI predicts individual differences in brain activity during task performance. Science (80-) 352: 216–220.

21. Finn ES, Todd Constable R (2016): Individual variation in functional brain connectivity: Implications for personalized approaches to psychiatric disease. Dialogues Clin Neurosci 18: 277–287.

22. Fox MD, Greicius M (2010): Clinical applications of resting state functional connectivity. Front Syst Neurosci 4. https://doi.org/10.3389/fnsys.2010.00019

23. Parker Jones O, Voets NL, Adcock JE, Stacey R, Jbabdi S (2017): Resting connectivity predicts task activation in pre-surgical populations. NeuroImage Clin 13: 378–385.

24. Kay SR, Fiszbein A, Opler LA (1987): The positive and negative syndrome scale (PANSS) for schizophrenia. Schizophr Bull 13: 261–276.

25. Owen AM, McMillan KM, Laird AR, Bullmore E (2005): N-back working memory paradigm: A meta-analysis of normative functional neuroimaging studies. Hum Brain Mapp 25: 46–59.

26. Livny A, Cohen K, Tik N, Tsarfaty G, Rosca P, Weinstein A (2018): The effects of synthetic cannabinoids (SCs) on brain structure and function. Eur Neuropsychopharmacol 28: 1047–1057.

27. Jansma JM, Ramsey NF, Van Der Wee NJA, Kahn RS (2004): Working memory capacity in schizophrenia: A parametric fMRI study. Schizophr Res 68: 159–171.

28. Krieger S, Lis S, Cetin T, Gallhofer B, Meyer-Lindenberg A (2005): Executive function and cognitive subprocesses in first-episode, drug-naive schizophrenia: An analysis of N-back performance. Am J Psychiatry 162: 1206–1208.

29. Smith SM, Jenkinson M, Woolrich MW, Beckmann CF, Behrens TEJ, Johansen-Berg H, et al. (2004): Advances in functional and structural MR image analysis and implementation as FSL. Neuroimage 23: S208–S219.

30. Griffanti L, Salimi-Khorshidi G, Beckmann CF, Auerbach EJ, Douaud G, Sexton CE, et al. (2014): ICA-based artefact removal and accelerated fMRI acquisition for improved resting state network imaging. Neuroimage 95: 232–247.

31. Glasser MF, Sotiropoulos SN, Wilson JA, Coalson TS, Fischl B, Andersson JL, et al. (2013): The minimal preprocessing pipelines for the Human Connectome Project. Neuroimage 80: 105–124.

32. Marcus DS, Harwell J, Olsen T, Hodge M, Glasser MF, Prior F, et al. (2011): Informatics and data mining tools and strategies for the human connectome project. Front Neuroinform 5: 1–12.

33. Woolrich MW, Ripley BD, Brady M, Smith SM (2001): Temporal autocorrelation in univariate linear modeling of FMRI data. Neuroimage 14: 1370–1386.

34. Woolrich MW, Behrens TEJ, Beckmann CF, Jenkinson M, Smith SM (2004): Multilevel linear modelling for FMRI group analysis using Bayesian inference. Neuroimage 21: 1732–1747.

35. Beckmann CF, Smith SM (2004): Probabilistic Independent Component Analysis for Functional Magnetic Resonance Imaging. IEEE Trans Med Imaging 23: 137–152.

36. Schaefer A, Kong R, Gordon EM, Laumann TO, Zuo X-N, Holmes AJ, et al. (2018): Local-Global Parcellation of the Human Cerebral Cortex from Intrinsic Functional Connectivity MRI. Cereb Cortex 28: 3095–3114.

37. Smith SM, Hyvärinen A, Varoquaux G, Miller KL, Beckmann CF (2014): Group-PCA for very large fMRI datasets. Neuroimage 101: 738–749.

38. Beckmann CF, DeLuca M, Devlin JT, Smith SM (2005): Investigations into resting-state connectivity using independent component analysis. Philos Trans R Soc B Biol Sci 360: 1001–1013.

39. Hyvärinen A (1999): Fast and robust fixed-point algorithms for independent component analysis. IEEE Trans Neural Networks 10: 626–634.

40. Breiman L (2001): Statistical Modeling: The Two Cultures (with comments and a rejoinder by the author). Stat Sci 16: 199–231.

41. Thomas Yeo BT, Krienen FM, Sepulcre J, Sabuncu MR, Lashkari D, Hollinshead M, et al. (2011): The organization of the human cerebral cortex estimated by intrinsic functional connectivity. J Neurophysiol 106: 1125–1165.

42. Saygin ZM, Osher DE, Koldewyn K, Reynolds G, Gabrieli JDEE, Saxe RR (2012): Anatomical connectivity patterns predict face selectivity in the fusiform gyrus. Nat Neurosci 15: 321–327.

43. Hampson M, Driesen NR, Skudlarski P, Gore JC, Constable RT (2006): Brain Connectivity Related to Working Memory Performance. 26: 13338–13343.

44. Cuthbert BN, Insel TR (2010): Toward New Approaches to Psychotic Disorders: The NIMH Research Domain Criteria Project. 36: 1061–1062.

45. Cetin MS, Calhoun VD, Miller R, Rashid B, Pearlson GD, Damaraju E, Arbabshirani MR (2016): Classification of schizophrenia and bipolar patients using static and dynamic resting-state fMRI brain connectivity. Neuroimage 134: 645–657.

46. Li A, Zalesky A, Yue W, Howes O, Yan H, Liu Y, et al. (2020): A neuroimaging biomarker for striatal dysfunction in schizophrenia. Nat Med. https://doi.org/10.1038/s41591-020-0793-8

47. Anderson A, Cohen MS (2013): Decreased small-world functional network connectivity and clustering across resting state networks in schizophrenia: An fMRI classification tutorial. Front Hum Neurosci 7: 1–18.

48. Hoptman MJ, Zuo XN, D’Angelo D, Mauro CJ, Butler PD, Milham MP, Javitt DC (2012): Decreased interhemispheric coordination in schizophrenia: A resting state fMRI study. Schizophr Res 141: 1–7.

49. Li HJ, Xu Y, Zhang KR, Hoptman MJ, Zuo XN (2015): Homotopic connectivity in drug-naïve, first-episode, early-onset schizophrenia. J Child Psychol Psychiatry Allied Discip 56: 432–443.

50. Guo W, Jiang J, Xiao C, Zhang Z, Zhang J, Yu L, et al. (2014): Decreased resting-state interhemispheric functional connectivity in unaffected siblings of schizophrenia patients. Schizophr Res 152: 170–175.

51. Eryilmaz H, Tanner AS, Ho NF, Nitenson AZ, Silverstein NJ, Petruzzi LJ, et al. (2016): Disrupted working memory circuitry in schizophrenia: Disentangling fMRI markers of core pathology vs other aspects of impaired performance. Neuropsychopharmacology 41: 2411–2420.

52. Dodds CM, Morein-Zamir S, Robbins TW (2011): Dissociating inhibition, attention, and response control in the frontoparietal network using functional magnetic resonance imaging. Cereb Cortex 21: 1155–1165.

