## Supplemental Information for "Predicting Individual Variability in Task-Evoked Brain Activity in Schizophrenia"

### Supplementary Materials


Niv Tik[1,2], Abigail Livny[1,3,4], Shachar Gal[1,2], Karny Gigi[5], Galia Tsarfaty[1,3], Mark Weiser[1,5] and Ido Tavor[1,2]

[1]Sackler Faculty of Medicine, Tel Aviv University, Tel Aviv, Israel

[2]Sagol School of Neuroscience, Tel Aviv University, Tel Aviv, Israel

[3]Division of Diagnostic Imaging, Sheba Medical Center, Tel-Hashomer, Israel

[4]Joseph Sagol Neuroscience Center, Sheba Medical Center, Tel-Hashomer, Israel

[5]Department of Psychiatry, Sheba Medical Center, Tel-Hashomer, Israel

### Schizophrenia (SCZ) Group Individual Characteristics

**Table S1.** SCZ individual demographics and clinical scores

| Age (years) | Gender | PANSS positive | PANSS negative | PANSS general | PANSS total |
|---|---|---|---|---|---|
| 24 | M | 7 | 9 | 15 | 31 |
| 26 | F | 20 | 12 | 28 | 60 |
| 35 | M | 24 | 25 | 34 | 83 |
| 32 | M | 10 | 21 | 24 | 55 |
| 32 | M | 9 | 18 | 25 | 52 |
| 30 | M | 8 | 10 | 19 | 37 |
| 28 | M | 7 | 9 | 24 | 40 |
| 40 | M | 10 | 19 | 21 | 50 |
| 20 | M | NA | NA | NA | NA |
| 39 | F | NA | NA | NA | NA |
| 21 | M | 8 | 13 | 20 | 41 |
| 24 | M | 13 | 13 | 25 | 51 |
| 30 | M | 7 | 12 | 18 | 37 |
| 40 | M | 18 | 15 | 23 | 56 |
| 24 | F | 10 | 21 | 17 | 48 |
| 19 | M | 10 | 23 | 32 | 65 |
| 30 | M | 7 | 16 | 18 | 41 |
| 31 | M | 9 | 16 | 25 | 50 |
| 26 | F | NA | NA | NA | NA |
| 35 | M | 10 | 23 | 31 | 64 |

**fMRI Working Memory Task**

The fMRI task all participants conducted was a version of the well-validated N-back task (1). In this task, a continuous sequence of stimuli is presented to the participants and they are asked to indicate whether the current stimulus matches a stimulus presented ‘n’ (1 or 2) stimuli earlier in the sequence. The specific design in this study used Hebrew letter stimuli presented at three different memory load conditions (0back ,1back, 2back) (2). In each condition Hebrew letters were presented on the screen and the participant was asked to press a button with the right index finger every time the stimulus matched the rule. The number of target stimuli was equal across all three conditions and appeared in 33% of the trials for each condition. Each stimulus was presented for 0.5S, followed by a 1.5S of a blank screen inter-stimulus interval. In the 0back condition the participant responded every time a target letter was presented (No memory load). In the 1back conditions the participant was instructed to respond each time a stimulus was presented in two consecutive trials (Low memory load). In the 2back condition the participant was instructed to respond every time the stimulus presented matched the one presented two trials before (High memory load). The task included Twelve blocks: six 0back blocks, three 1back blocks and three 2back blocks. Participants that failed to reach 70% correct answers in one or more of the three conditions were excluded from the study. See task description in Figure S1.

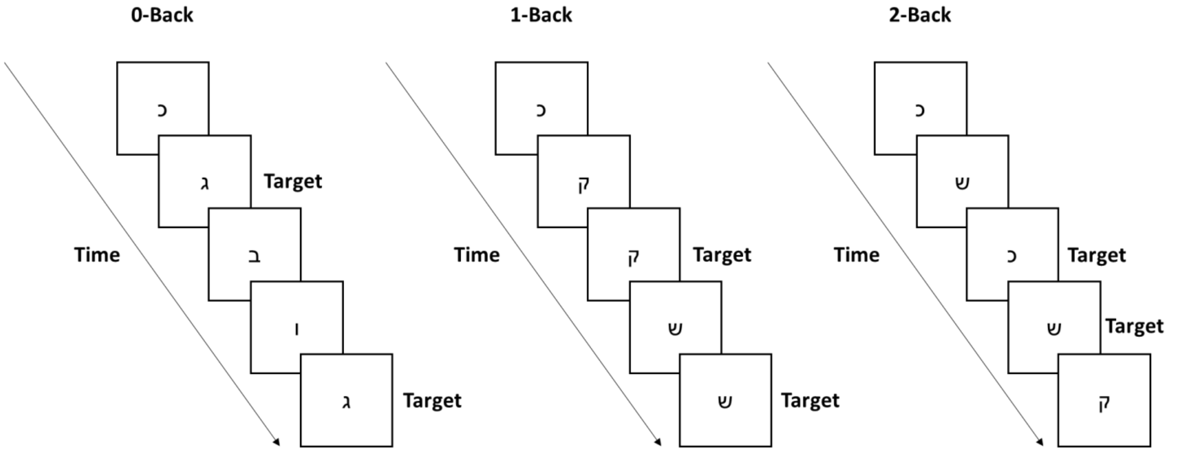


**Figure S1.** N-back working memory task design. In each condition, participants pressed a button with their right index finger every time a target stimulus appeared on the screen. Each stimulus was presented for 0.5S followed by a 1.5S blank screen interstimulus interval. Total of twelve blocks were presented: six 0back, three 1back and three 2back blocks.

### Functional Connectivity-based Classification

For classification, we used an elastic-net logistic regression model, that combines L1 and L2 penalties for regularization. The L1/L2 ratio ($\alpha$) and the regularization factor ($\lambda$) were chosen in a stratified 5-fold cross-validation procedure. Each fold contained 4 SCZ patients and 16 controls. ROC (receiver operating characteristic) curve for the 5-fold cross-validation is depicted in Figure S2.

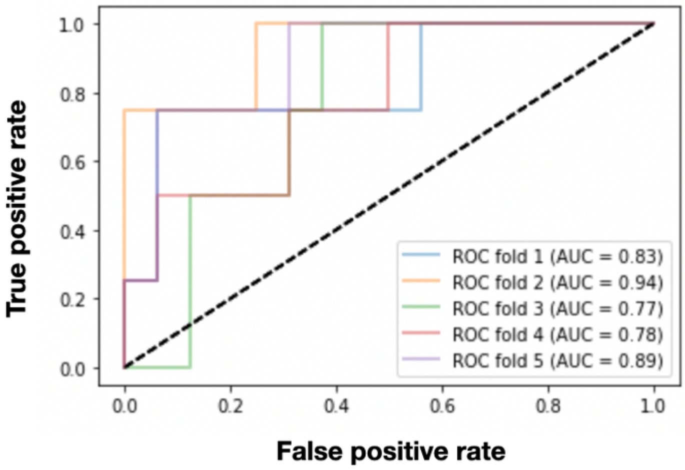


**Figure S2.** ROC curve showing False positive rate (FPR, X-axis) and True positive rate (TPR, Y-axis) across a 5-fold cross validation. Calculated area under the curve (AUC) for each fold is presented on the legend. Dashed line represents equality between FPR and TPR (AUC=0.5) which is what is expected under random classification

In order to determine chance level rates and utilize them to test for significance we used a 5000 iterations permutation test. We used a split-half routine for this permutation test: half of the data was used to train the model that will then be used to classify the other half. Then, we performed 5000 iterations in which we shuffled the labels of the test set randomly and calculated chance levels for 4 different scores: AUC (area under the ROC curve), accuracy, sensitivity and specificity by comparing the predictions to the shuffled labels.

The scores were calculated as such:

$$\text{Accuracy} = \frac{true\ positive + true\ negative}{true\ positive + true\ negative + false\ positive + false\ negative}$$

$$\text{Sensitivity} = \frac{true\ positive}{true\ positive + false\ negative}$$

$$\text{Specificity} = \frac{true\ negative}{true\ negative + false\ positive}$$

AUC: Average area under the receiver operating characteristics (ROC) curve across all cross-validation folds.

The significance of each score was determined as such:

$$\frac{number\ of\ values\ equal\ or\ higher\ to\ the\ value\ calculated\ using\ the\ true\ labels}{number\ of\ iterations}$$

In order to examine the characteristics of the features that contributed most to the prediction model, we trained the classifier on all the data, and sorted the features according to their absolute beta values. It is of importance to note that the proportion of homotopic connections in the whole data set was only 8%, whereas its proportion in the top 100 features was much larger (31%), and 70% in the top 10 (Figure S3). To further explore the contribution of homotopic connections to the classification model, we performed the classification based on homotopic edges only (Figure S4), which yielded improved classification scores (Table S2).

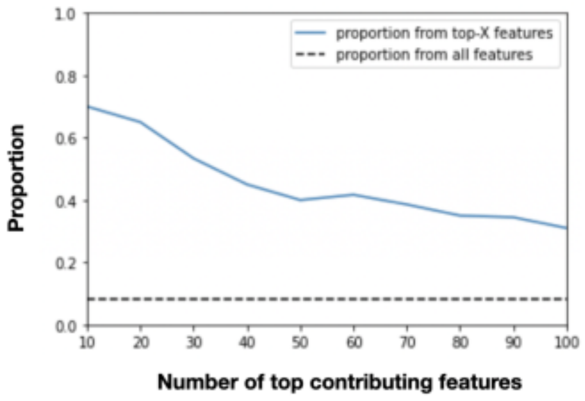


**Figure S3** Proportion of homotopic edges (Y-axis) out of the number of top contributing features (X-axis) in our classification model, which is remarkably higher than the proportion of homotopic connections in all the features that were used for classification (marked by a dashed horizontal line).

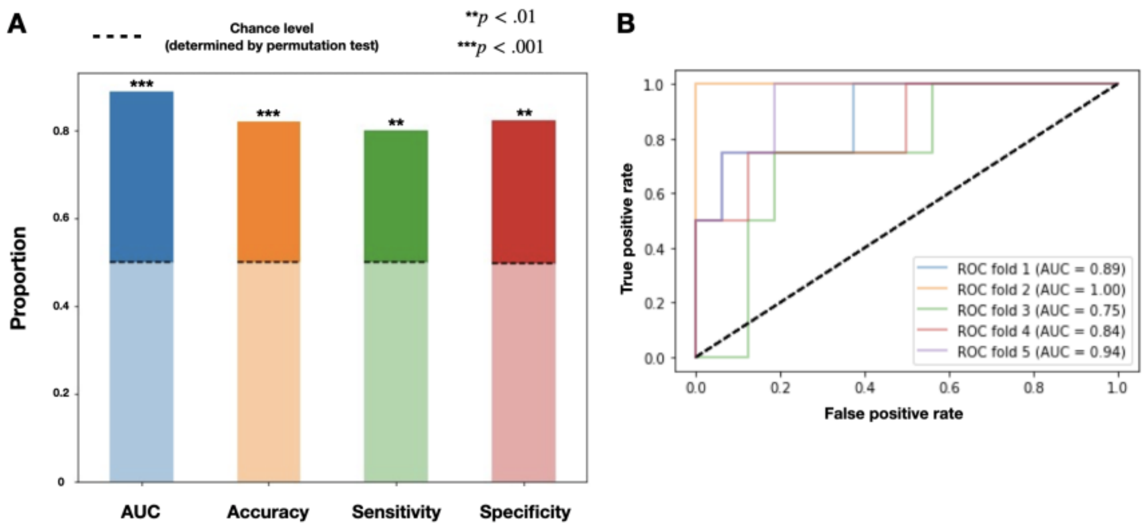


**Figure S4.** Classification using homotopic edges only. (A) shows the results of a permutation test designed to determine classification success. We show a bar graph with 4 scores: area under the ROC curve (AUC), accuracy, sensitivity and specificity. Computed chance level for each score is marked by a dashed line on the bars. (B) displays a ROC curve showing False positive rate (FPR, X-axis) and True positive rate (TPR, Y-axis) across a 5-fold cross validation using homotopic edges only. Calculated area under the curve (AUC) for each fold is presented on the legend. Dashed line represents equality between FPR and TPR (AUC=0.5) which is what is expected under random classification.

**Table S2.** Comparison between a classification using features from the entire cortex to one using homotopic features only. The score presented were calculated using train-test split, and the p-values were obtained using a 5000 iterations permutation test. The higher value for each performance score is highlighted, indicating equal or improved performance for the homotopic only classification in all calculated measures

| | **All cortex features** | **Homotopic features only** |
|---|---|---|
| **AUC** | 0.85 ($p = 0.0002$) | **0.89** ($p = 0.0002$) |
| **Accuracy** | 0.78 ($p = 0.003$) | **0.82** ($p = 0.0008$) |
| **Sensitivity** | **0.8** ($p = 0.003$) | **0.8** ($p = 0.0038$) |
| **Specificity** | 0.77 ($p = 0.0074$) | **0.825** ($p = 0.0018$) |

### Prediction Pipeline

The first step of the prediction pipeline was aimed to extract features that will be used as predictors for the model from the preprocessed surface represented rs-fMRI scans. The group level steps were performed only on the training set control group. Resting-state fMRI data were processed using group-PCA (3), reducing dimensionality to a set of 200 principal components. Following, group-ICA (4) for all cortical components was performed using fast ICA (5), yielding 60 cortical connectivity maps that were used as features for our GLM model.

We used dual regression against individual time series to produce subject-specific cortical ICA maps (6) for both control and SCZ groups. The first regression used the cortical group maps described above as regressors to get individual time series for each subject and each map. The second regression used the individual time series as regressors to get individual spatial maps. We focused on predicting task data on the cortex, which is where we observe the majority of the inter-subject variability.

A general linear model was used to map the functional connectivity features to the task data (i.e. individual z-score contrast maps derived from fMRI task analysis). The task data were modelled using the general linear model, broken down spatially into 50 non-overlapping regions of interest created using group ICA and a winner-takes-all parcellation on the resulting ICA maps (based on Tavor et al., 2016 (7)). Within each of these 50 parcels, a general linear model was used to fit features to the activation data in each participant. Prior to learning the model parameters, all features were normalized to zero mean and unit norm across the whole brain. Feature normalization is important both to average regression results across participants and to compare regression coefficients between features. A constant regressor was added and the model fitted for each of the 50 parcels. To cover the whole cortex, we thus fitted 50 models for each subject. To generate predictions in our training set, we used a leave-one-out routine. Our model learned the regression coefficients for each subject and each brain parcel separately, the coefficients were averaged across $n - 1$ training set control participants for out-of-sample predictions. To predict an individual's task activation map, regression coefficients were averaged across all other subjects. This process was repeated for all 80 participants in the training set. Following, we used a model based on all 80 training set participants to predict brain activations in the SCZ test set.

### References


1. Owen AM, McMillan KM, Laird AR, Bullmore E (2005): N-back working memory paradigm: A meta-analysis of normative functional neuroimaging studies. *Hum Brain Mapp* 25: 46–59.

2. Livny A, Cohen K, Tik N, Tsarfaty G, Rosca P, Weinstein A (2018): The effects of synthetic cannabinoids (SCs) on brain structure and function. *Eur Neuropsychopharmacol* 28: 1047–1057.

3. Smith SM, Hyvärinen A, Varoquaux G, Miller KL, Beckmann CF (2014): Group-PCA for very large fMRI datasets. *Neuroimage* 101: 738–749.

4. Beckmann CF, DeLuca M, Devlin JT, Smith SM (2005): Investigations into resting-state connectivity using independent component analysis. *Philos Trans R Soc B Biol Sci* 360: 1001–1013.

5. Hyvärinen A (1999): Fast and robust fixed-point algorithms for independent component analysis. *IEEE Trans Neural Networks* 10: 626–634.

6. Breiman L (2001): Statistical Modeling: The Two Cultures (with comments and a rejoinder by the author). *Stat Sci* 16: 199–231.

7. Tavor I, Jones OP, Mars RB, Smith SM, Behrens TE, Jbabdi S (2016): Task-free MRI predicts individual differences in brain activity during task performance. *Science (80- )* 352: 216–220.
